# Maximum Classifier Discrepancy Generative Adversarial Network for Jointly Harmonizing Scanner Effects and Improving Reproducibility of Downstream Tasks

**DOI:** 10.1101/2022.11.19.517154

**Authors:** Weizheng Yan, Zening Fu, Rongtao Jiang, Jing Sui, Vince D. Calhoun

## Abstract

**Objective:** Multi-site collaboration is essential for overcoming small-sample problems when exploring reproducible biomarkers in MRI studies. However, various scanner-specific factors dramatically reduce the cross-scanner replicability. Moreover, existing harmony methods mostly could not guarantee the improved performance of downstream tasks.

**Methods:** we proposed a new multi-scanner harmony framework, called ‘maximum classifier discrepancy generative adversarial network’, or MCD-GAN, for removing scanner effects in the original feature space while preserving substantial biological information for downstream tasks. Specifically, the adversarial generative network was utilized for persisting the structural layout of each sample, and the maximum classifier discrepancy module was introduced for regulating GAN generators by incorporating the downstream tasks.

**Results:** We compared the MCD-GAN with other state-of-the-art data harmony approaches (*e*.*g*., ComBat, CycleGAN) on simulated data and the Adolescent Brain Cognitive Development (ABCD) dataset. Results demonstrate that MCD-GAN outperformed other approaches in improving cross-scanner classification performance while preserving the anatomical layout of the original images.

**Significance:** To the best of our knowledge, the proposed MCD-GAN is the first generative model which incorporates downstream tasks while harmonizing, and is a promising solution for facilitating cross-site reproducibility in various tasks such as classification and regression. The codes of the MCD-GAN are available at https://github.com/trendscenter/MCD-GAN.

## I. Introduction

Multi-site neuroimaging collaboration is a viable way to overcome small-sample biases by aggregating samples from multiple sites or hospitals. However, samples from different sites are usually collected using various scanning manufacturers, acquisition protocols, and software versions. As illustrated in Fig. 1, the extracted cortical thickness feature from distinct MRI scanners displays significant group differences, leading to a loss of efficiency when applying the trained models or discovered biomarkers from one scanner to the other. This partly explains the reason for significant deterioration in pooled classification performance as the sample size increases [1]. Hence, properly harmonizing site/scanner effects is critical for improving the outcomes of large-sample studies [2].

**Fig. 1.**
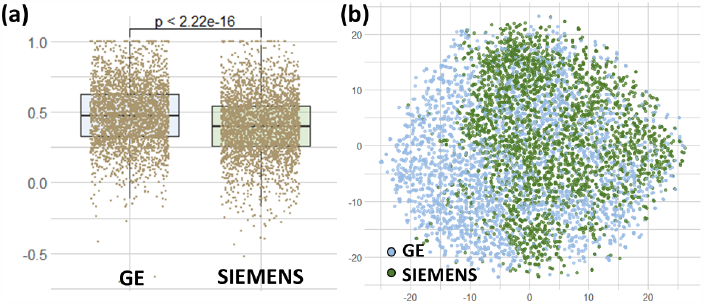
Visualization of scanner effects. (a) Comparison of the cortical thickness of a specific brain region (left hemisphere supramarginal ROI, z-scored) scanned by GE Discovery (n=2860 subjects) and SIEMENS Prisma (n=3075); (b) TSNE visualization of all cortical thickness features (n=68) of all subjects collected using GE Discovery (n=2860) and SIEMENS Prisma (n=3075). *Notes*: The cortical thickness features were preprocessed by the ABCD research consortium and are downloaded from https://nda.nih.gov/abcd/.

Non-biological confounds often have unpredictable prior distributions, making it challenging to be modeled. Most of the existing image harmonization algorithms [3] can be categorized into two types: residual-based [4-8] and generative adversarial network (GAN)-based methods [2, 9-11]. Residual-based approaches (*e*.*g*., residual regression, ComBat [5, 7], and Neuroharmony [4]) harmonize samples from different scanners by estimating the distribution of confounds based on three assumptions: *1)* independence: the mutual correlations between features can be ignored; *2)* linearity: the distribution of the confounds can be estimated and removed using linear models; *3)* simultaneity: all samples to be harmonized are accessible at the same time. ComBat and its variants perform a Bayesian regression that corrects the measurements from different samples with addictive and multiplicative terms, and have been successfully applied for harmonizing multi-site cortical thickness features especially when the sample size is relatively small. However, ComBat requires all samples scanned by various scanners to be put together for harmonization, which means the analysis finished on the harmonized dataset should be re-run if new samples scanned using different scanners are to be added, leading to a considerable amount of repetitive work.

Compared to residual-based approaches which usually ignore the mutual correlations among features, GAN-based approaches can capture more complex non-linear relations using multiple layers [12]. Cycle-consistent generative adversarial network (CycleGAN), has been applied for multi-site MRI harmonization [10]. Another advantage of GAN-based approaches is they do not require samples of all scanners to be accessed at the same time. Instead, they use samples from a specific scanner to build the source domain, then map samples from other scanners to the source domain, making continuous harmonization possible.

However, neither residual-based nor GAN-based harmony approaches can guarantee improved performances for specific downstream tasks. Even worse, if the scanner confounds are estimated incorrectly, harmonizing may even degrade the performance of downstream tasks. To overcome this issue, the downstream task should be incorporated when harmonizing. Domain adaptation [13] aims to theoretically guarantee that the model trained on the source domain can also achieve high performance on the target domain. The domain adaptation approach usually trains a task-specific model by mapping both source and target samples into a domain-shared subspace. Maximum classifier discrepancy (MCD) [14], which utilizes the discrepancy of two classifiers for constraining mappings from original space to a domain-shared subspace, is a practical framework in the domain adaptation field. However, the original MCD implementation could not maintain anatomical information, limiting its application in the medical imaging field.

In this work, by combining the advantages of both harmonization and domain adaptation, we aimed to design a new data harmony method named maximum classifier discrepancy generative adversarial network (MCD-GAN). The MCD-GAN has three strengths as follows: *1)* harmonizing multi-scanner samples while preserving their anatomical layouts; *2)* incorporating the downstream task information to modulate the harmonization for improving the downstream task performance; *3)* compatibility for continuous harmonization.

The paper is organized as follows. Section II describes the related data harmony and domain adaptation methods including residual-based, CycleGAN, and Maximum Classifier Discrepancy. Second III describes the methodological overview, network architecture, learning objectives, training steps, and theoretical analysis of the proposed MCD-GAN. In section IV, the MCD-GAN is applied to simulated data and the Adolescent Brain Cognitive Development (ABCD) dataset. The harmonization and classification are compared with ComBat and CycleGAN. In section V, the advantages, limitations, and future directions are further discussed. The MCD-GAN code is available via https://github.com/trendscenter/MCD-GAN.

## II. Related Work

### Residual-based harmony

Residual-based approaches typically harmonize data using linear regression. ComBat is a method adopted from the genomic literature [8], and has been applied for harmonizing diffusion tensor imaging [7] and cortical thickness [5] in multiple sites samples. ComBat extends the conventional residual regression by modeling site-specific scaling factors and using empirical Bayesian to improve the estimation of site parameters for small sample sizes. The model assumes that the expected value of the imaging features can be modeled as a linear combination of biological variables and scanner effects, whose error term is modulated by additional scanner-specific scaling factors. The ComBat can remove unwanted confounds associated with the scanner while preserving other biological associations in the samples.

### CycleGAN harmony

CycleGAN utilizes two generators, *G*_*s*→*t*_ and *G*_*t*→*s*_, to learn the the mappings from source domain to target domain and its inverse mapping. Two discriminators, *D*_*s*_ and *D*_*t*_, utilize adversarial loss measure how realistic the generated images (*G*_*s*→*t*_ (*X*_*s*_) ≈ *X*_*t*_ or *G*_*t*→*s*_ (*X*_*t*_) ≈ *X*_*s*_) are by an adversarial loss and how well the original input is reconstructed after a sequence of two generators (*G*_*s*→*t*_(*G*_*t*→*s*_ (*x*_*t*_)) ≈ *x*_*t*_ or *G*_*t*→*s*_(*G*_*s*→*t*_ (*x*_*s*_)) ≈ *x*_*s*_)) by circle consistency losses. Thus, the objective of training the CylceGAN is to make the distribution of images generated from *G*_*s*→*t*_ (*x*_*s*_) (or *G*_*t*→*s*_ (*x*_*t*_)) indistinguishable from the distribution *x*_*t*_ (or *x*_*s*_).

CycleGAN has been applied to MRI for multi-site harmony while preserving the anatomical layout of the MRI [9, 10, 15]. For example, Bashyam et al utilized a modified CycleGAN architecture for removing site effects and achieved improved cross-site age prediction performance than without harmonization.

### Maximum Classifier Discrepancy (MCD)

MCD is a method proposed for aligning distributions of source and target by utilizing task-specific decision boundaries [14, 16]. MCD consists of two core modules: two task-specific classifiers and one domain-shared feature extractor. The two task-specific classifiers are trained for the specific downstream task (*e*.*g*., image classification). The classifiers are optimized to identify the category of source domain samples by taking features from the feature extractors. When applying the trained classifiers to target domain samples, some samples in the target domain which are far from the support of the source domain are misclassified. By maximizing the discrepancy between the two classifiers, the target domain samples which are far from the support of the source domain can be detected. As for the domain-shared feature extractor, it is optimized for generating target features for minimizing the discrepancy between two classifiers. In this way, the feature extractor is constrained to generating features that have more consistency between source and target domains. One problem of the MCD approach is it maps features to a subspace without preserving its semantical information, limiting its usage in neuroimaging studies. A comprehensive comparison of the above algorithms is shown in Table 1.

**Table 1.** Comparisonof site-effects harmony approaches.

|  | Continuous harmony? | Task-specific? | Non-linear? | Preserve layout? |
| --- | --- | --- | --- | --- |
| Combat [5] | × | × | × | ✓ |
| Cycle-GAN [10] | ✓ | × | ✓ | ✓ |
| MCD [14] | × | ✓ | ✓ | × |
| MCD-GAN | ✓ | ✓ | ✓ | ✓ |

## III. Methods

### A Methodological Overview

Fig.2. is a conceptual overview of our proposed MCD-GAN framework consisting of two core modules: CycleGAN and two classifiers. The CycleGAN contained two generators (*G*_*s*→*t*_ and *G*_*t*→*s*_), and respective adversarial discriminators (*D*_*s*_ and *D*_*t*_). *D*_*s*_ constrained *G*_*s*→*t*_ to generate high-quality samples from the source domain to the target domain. *D*_*t*_ constrained *G*_*s*→*t*_ to generate samples from the target domain to the source domain. The two classifiers (*F*_1_ and *F*_2_) were first optimized for classifying source domain samples. The optimized classifiers were then applied to the unlabeled generated samples from the target domain to get predicted labels respectively. The discrepancy between the two classifiers was obtained by comparing the different classification results of the two classifiers on generated samples. Two steps as follows were iteratively conducted to max-min the classifier discrepancy: ***1)*** Optimize the two classifiers (*F*_1_ and *F*_2_) to maximize the discrepancy on ‘fake’ (generated from the target domain using *G*_*t*→*s*_) samples while maintaining the classification performance on source domain samples; ***2)*** Optimize the *G*_*t*→*s*_ for minimizing the classification discrepancy on ‘fake’ samples.

**Fig. 2.**
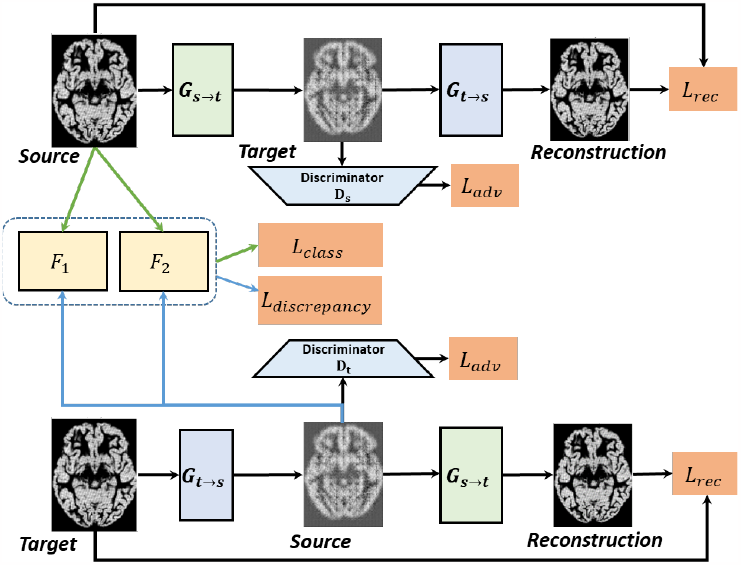
Overview of the proposed MCD-GAN model. The model contains two generative modules, *G*_*s*→*t*_ and *G*_*t*→*s*_, and corresponding adversarial discriminators, *D*_*s*_ and *D*_*t*_. Through adversarial training, the *G*_*s*→*t*_ can learn to map source domain samples into the target domain, and the *G*_*t*→*s*_ can learn to map target domain samples into the source domain. Two classifiers, *F*_1_ and *F*_2_, are optimized for maximizing the discrepancy of generated source sample’s classification results (shown as blue pipelines) while maintaining the classification performance on source domain samples (shown as green pipelines). The *G*_*t*→*s*_ is also optimized according to the discrepancy for regulating the mapping from the target domain to the source domain.

### B Model Architecture and Design Details

The proposed MCD-GAN model is a general framework consisting of generators, discriminators, and classifiers. The details of the architecture can be modified to fit the characteristics of the input features. In this study, three datasets (“double moon” (Fig. 3a), cortical thickness vectors (Fig. 3b), and T1-weighted MR images (Fig. 3c)) were used for testing the performance of the MCD-GAN. Detailed architectures of the model for each dataset are shown below:

**Fig. 3.**
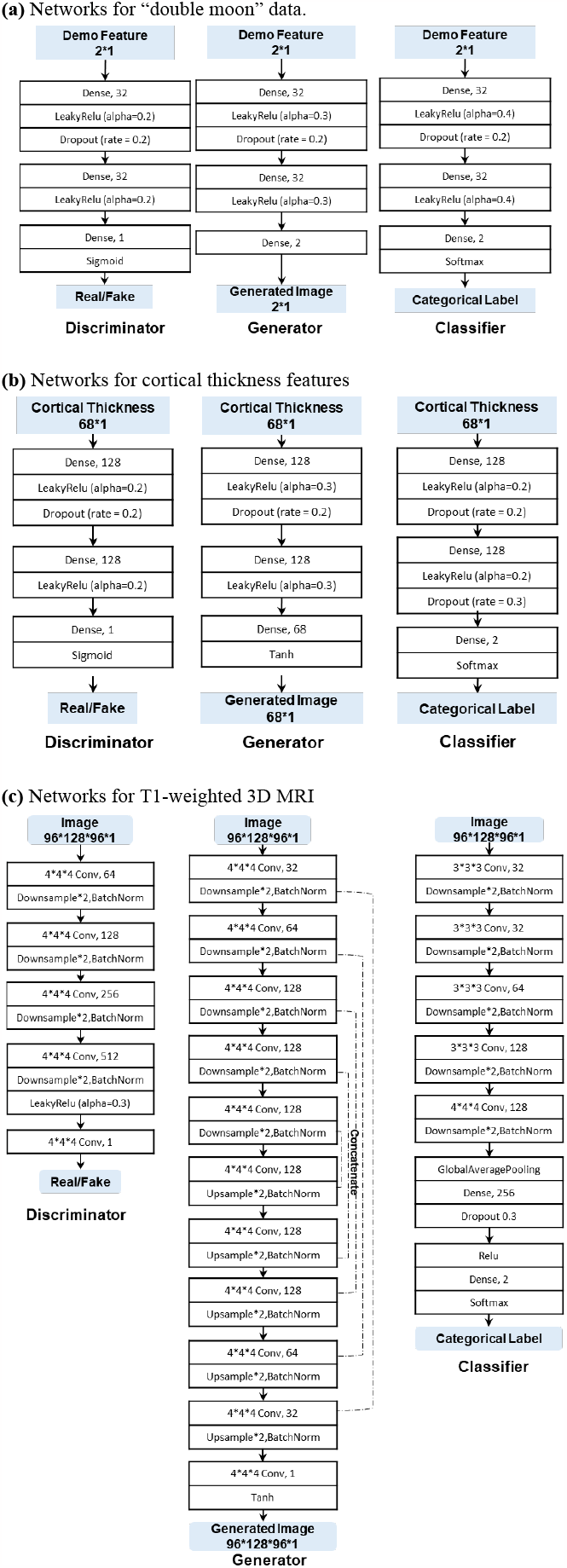
The architectures of the proposed MCD-GAN for different feature types. (a) Networks for “double moon”; (b) Networks for cortical thickness features; (c) Networks for T1-weighted MR images. Notes: from left to right are discriminators, generators, and classifiers.

#### 1) Generators

For “double moon” and cortical thickness vectors, fully connected networks are used as generators. For T1-weighted MR images, U-Nets are used as generators.

#### 2) Discriminators

For “double moon” and cortical thickness vectors, fully connected networks are used as the discriminators. For T1-weighted MR image features, U-Nets are used as the discriminators.

#### 3) Classifiers

For “double moon” and cortical features, fully connected networks are used as classifiers. For the T1-weighted MR image features, 3D-CNNs are used as classifiers.

### C Learning Objectives

The loss functions of the proposed MCD-GAN consist of adversarial loss, cycle-consistency loss, classification loss, and max-discrepancy loss. Details of the loss functions are listed as follows:

#### 1) Adversarial loss

The bidirectional cycles are utilized for performing global domain alignment in the adaptation process. The source domain loss *L*_*GAN*(_(*G*_*t*→*s*_, *D*_*s*_) and target domain loss *L*_*GAN*_ (*G*_*s*→*t*_, *D*_*t*_) are:

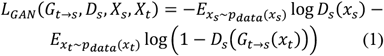

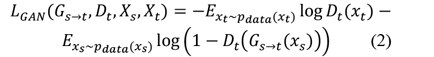

where *D*_*s*_ and *D*_*t*_ are discriminators corresponding to the source and target domains. *G*_*s*→*t*_ is the generator mapping source domain features to the target domain, *G*_*t*→*s*_ is the generator to map target domain features to the source domain. *X*_*s*_ and *X*_*t*_ are samples from the source domain and target domain respectively.

#### 2) Cycle-consistency loss

The cycle consistency loss is utilized to regularize the two generators (*G*_*s*→*t*_ and *G*_*t*→*s*_). The loss of cycle consistency is as follows:

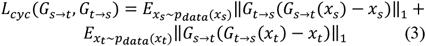

The CycleGAN loss is a weighted summation of the adversarial loss and cycle-consistency loss as follows:

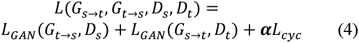

where *α* is the hyperparameter that controls the ratio between adversarial loss and cycle-consistency loss *α* is set to 10 in default [12].

#### 3) Classification loss

The classifiers are trained on source domain samples. The loss function is as follows:

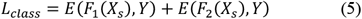

where *E* denotes the cross-entropy loss. *X*_*s*_ represent the samples from the source domain. *Y* represents the ground truth labels of samples. *F*_1_ and *F*_2_ are classifiers.

#### 4) Classifiers discrepancy loss

The two deep learning classifiers are first trained using samples from the source domain, then directly applied to samples generated from the target domain samples for testing and calculating the two classifiers’ discrepancy. Similar to [14], we utilize the absolute value of the difference between two classifiers’ probabilistic outputs as discrepancy loss:

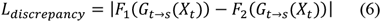

### D MCD-GAN training steps

The training steps of the proposed MCD-GAN are as follows:

#### Step A

The generators and discriminators are pre-trained from random to roughly align the source domain and target domain. The objective function is as follows:

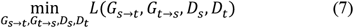

#### Step B

Two classifiers (*F*_1_ and *F*_2_) are trained using samples from the source domain. The optimized classifiers are then tested on the unlabeled ‘fake’ samples generated from the target domain samples using *G*_*t*→*s*_. The discrepancy between the two classifiers on unlabeled ‘fake’ samples is maximized by optimizing the classifiers while preserving the classification performance on the source domain. Therefore, the objective function in step B is as follows:

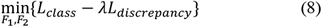

where λ is a hyperparameter that controls the ratio between the classification loss and the discrepancy loss. The effect of the λ is further discussed in the “discussion” section.

#### Step C

The generator*G*_*t*→*s*_ is optimized to minimize the discrepancy between the two classifiers. The term denotes the trade-off between the generator and classifiers. The objective is as follows:

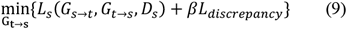

For simplicity, the *β* in (9) is set the same as the *λ* in (8). The above three steps are repeated until convergence.

### E Theoretical Analysis

The MCD-GAN is motivated by the domain adaptation theory proposed by Saito et al [14] and Ben-David et al [17]. Here, we introduce the relations between the MCD-GAN and the previous work. Ben-David et al [17]. proposed the theory that bounds the expected error on the target samples, *R*_*t*_(*h*), by using three terms: *1*) expected error on the source domain, *R*_*s*_(*h*); *2)* the discrepancy between two classifiers, *d*_*H*Δ*H*_(*S, T*), where *S* and *T* denote source and target domain respectively; 3) the shared error of the ideal joint hypothesis *ε*. Another theory proposed by Ben-Divid et al [18] bounds the error on the target domain and introduced domain divergence *d*_*H*_(*S, T*). The two theories and their relationships are explained as follows:

#### Theorem 1

Let *H* be the hypothesis class. Given two domains *S* and *T*, we have:

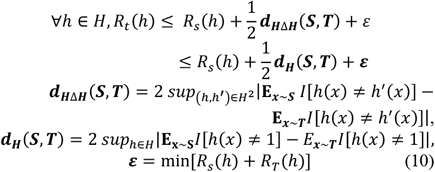

where *R*_*T*_(*h*) is the error of hypothesis *h* on the target domain, and *R*_*s*_(*h*) is the corresponding error on the source domain. *I*[a] is the indicator function, which is 1 if the predicate is true and 0 if false.

The *H* distance is empirically measured by the error of the domain classifiers which are trained to discriminate the domain of features. *ε* is a constant that is considered sufficiently low to achieve an accurate adaptation. Here, we further show the relationship between our MCD-GAN model and the *H*Δ*H* distance.

As for *d*_*H*Δ*H*_(*S, T*), if both the two classifiers (*h* and *h*^*’*^) can accurately classify source samples, the term E *I*_*x∼s*_[*h*(*x*) ≠ *h*^*’*^(*x*)] can be very small. *h* and *h*^*’*^ should be consistent on source samples. Thus, *d*_*H*Δ*H*_(*S, T*) will approximate the upper boundary of the expected disagreement of the two classifiers’ predictions on target samples.

We assume that *h* and *h*^*’*^ take features from a shared feature extraction module. Then we decompose the hypothesis *h* into *G*_*t*→*s*_ → *F*_1_, and *h*^*’*^ into *G*_*t*→*s*_ → *F*_2_, *G*_*t*→*s*_, *F*_1_ and *F*_2_ correspond to the network in our proposed MCD-GAN model. If we substitute those notations into the 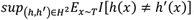, for fixed *G*_*t*→*s*_, the term will become 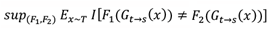. Furthermore, if we replace *sup* with max and minimize the term related to *G*_*t*→*s*_, we obtain

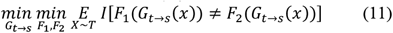

This is equivalent to the min-max problem we solve in MCD-GAN, in which classifiers are trained to maximize their discrepancy on target samples and the generator tries to minimize it. Therefore, the MCD-GAN which combines the MCD and CycleGAN is the framework that can not only maintain the anatomical layout of the samples but also improve the classification performance of downstream tasks.

## IV. Experiments

### A Data and Preprocessing

#### 1) Simulated data (double moon)

As shown in Fig. 4b. The double moon source domain (domain 1) data is simulated using the ‘*make_moons*’ function in Sklearn (https://scikit-learn.org/stable/). The noise hyper-parameter was set as 0.1. The number of source domain samples was set as 1000. The samples at the top ‘moon’ belong to category 1 and the samples at the bottom ‘moon’ belong to category 2. By rotating the source domain 30 degrees counterclockwise, we obtained target domain (domain 2) samples.

**Fig. 4.**
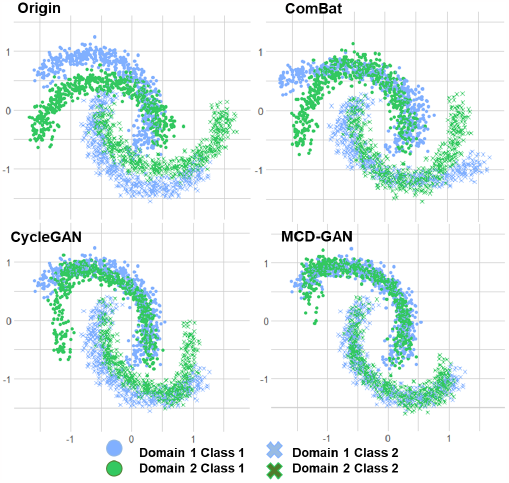
Harmonization comparison on simulated “double moon” dataset. Two domains (domain 1 and domain 2) of samples are simulated. Each domain consists of two categories (class 1 and class 2). Domain 2 is obtained by rotating domain 1 by 30 degrees counterclockwise. ***Notes***: top left: original samples; top right: samples after harmonizing using ComBat; bottom left: the samples of domain 2 are mapped to domain 1 using CycleGAN; bottom right: the samples of domain 2 are mapped to domain 1 using MCD-GAN.

#### 2) ABCD Cortical Thickness features

The ABCD study (https://abcdstudy.org) is a longitudinal, large-scale research project which collected multiple data types from over ten thousand US children to identify the internal and external factors that can affect an individual’s developmental trajectory [19]. The data are available to qualified researchers via a repository managed by the National Institute of Mental Health Data Archive (NDA; https://nda.nih.gov/abcd). The data were collected using scanners of three main manufacturers including GE, SIEMENS, and Philips. In this work, the samples scanned using GE Discovery and SIEMENS Prisma are selected for analysis. The cortical thickness features are preprocessed as in [20]. The demographic information is shown in Table 2.

**Table 2.** Demographic information of the abcd dataset.

|  | GE<br>(Discovery) | SIEMENS<br>(Prisma) |
| --- | --- | --- |
| Cortical thickness (68 features) | 2860 subjects | 3075subjects |
| sMRI (121*145*121 voxels) | 2860 subjects | 3075subjects |
| Gender(F/M) | 1376/1484 | 1433/1642 |
| Age of months(mean±std) | 118.2±7.6 | 119.3±7.5 |

#### 3) ABCD T1-weighted MR images

T1-weighted MR images scanned using GE Discovery and SIEMENS scanners are included. The MRI data were segmented into tissue probability maps for gray matter, white matter, and cerebral spinal fluid using SPM12 toolbox. The gray matter images were then warped to standard space, modulated, and smoothed using a Gaussian kernel with an FWHM = 10 mm. The preprocessed gray matter volume images have a dimensionality of 121 × 145 × 121 in the voxel space, with a voxel size of 1.5 × 1.5 × 1.5 mm^3^. The demographic information is shown in Table 2.

### B Implementation Details

The proposed models were implemented based on Tensorflow2 (https://www.tensorflow.org/). Adam with an initial learning rate of 10^−4^ was used as the optimizer for all models. All the above models were implemented on the cluster (Intel(R) Xeon(R) Gold 6230 CPU @ 2.10GHz, 20 CPU cores) with a GPU (Tesla V100-SXM2-32GB). ComBat was implemented based on the codes downloaded from https://github.com/Jfortin1/ComBatHarmonization. The detailed architectures of CycleGAN and MCD-GAN are described in Fig. 3. More details of the model can be found in our codes https://github.com/trendscenter/MCD-GAN.

### C Simulated data results

As shown in Fig. 4, the performance of the proposed MCD-GAN was compared with ComBat and CycleGAN on simulated ‘double moon’ data. Before harmony, the classifier trained on domain 1 could not perform well on domain 2 (fig. 4, top left) because of the inconsistency of the two domains. The ComBat harmonized the two domains by rescaling and relocating samples in both two domains. However, ComBat was not able to model or remove the nonlinear confounds such as the rotation effects. The Cycle-GAN could accurately model the non-linear site effects and achieved better performance than ComBat. The flaw of CycleGAN was it only modeled the global distribution of two domains without incorporating the downstream task information. As shown in the bottom left of Fig.4, the samples in domain 2 which were far from the center could not be correctly harmonized. Differently, MCD-GAN utilizes the downstream classification task information as a constraint and therefore achieved the best performance. In addition, as can be seen in Fig. 4, the ComBat required the change of both source and target domain samples for harmony, however, the CycleGAN and MCD-GAN harmonized samples by mapping the domain 2 samples to domain 1 without changing the original domain 1 samples. Therefore, the GAN-based approaches are more suitable for continuous harmonizing when tackling multiple domains.

### D ABCD cortical thickness feature harmony

To compare how each data harmony algorithm changes the data, the numerical values of ABCD cortical thickness features are visualized. As shown in Fig. 5., before harmonization, the samples collected using GE discovery and SIEMENS Prisma exhibits significant differences. The ComBat harmonized the features by changing both the source domain (GE Discovery) and the target domain (SIEMENS Prisma). The CycleGAN and MCD-GAN keep the source domain (GE Discovery) unchanged while mapping the features in the target domain (SIEMENS Prisma) to the source domain (GE Discovery). In addition, we utilized tSNE for visualizing all ABCD samples (n=5935) by taking the cortical thickness features (n=68) as input. Before harmonizing, the samples from GE Discovery and SIEMENS Prisma exhibit clear cluster boundaries. All the harmony methods, including ComBat, CycleGAN, and MCD-GAN, could remove the scanner effects.

**Fig. 5.**
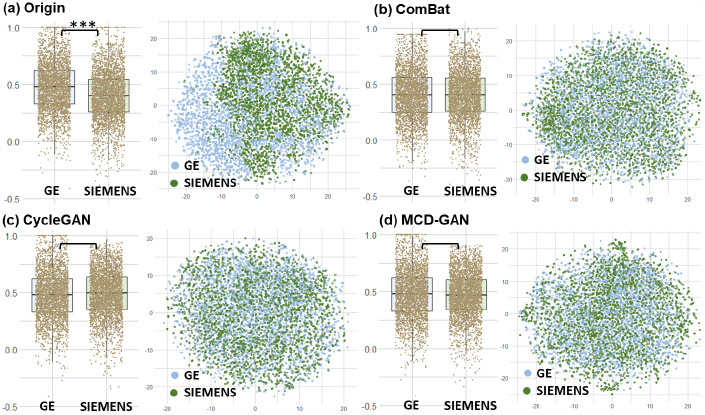
Harmonization comparison on ABCD cortical thickness features. Notes: The left figure in each subplot is the visualization of one specific cortical thickness feature (left hemisphere supramarginal ROI) scanned by GE Discovery and SIEMENS Prisma; the right figure in each subplot is the visualization of all cortical thickness features. ***p<0.001.

As shown in Figure 5b, ComBat harmonized scanner effects by changing the numeric value of all scanners. Differently, as shown in Figure 5c-d, GAN-based approaches mapped SIEMENS samples to GE domain for harmony. Therefore, assuming there are new samples scanned using Philips scanner to be harmonized, the GAN-based methods only need to map the Philips samples to GE domain. Therefore, compared to ComBat, the GAN-based methods are more flexible for continuous multi-scanner harmonizing especially when not all scanner samples are accessible at the same time.

### E Classification results after harmonization

The downstream cross-domain classification performances are compared with ComBat and CycleGAN. For the simulated ‘double moon’ data, the position of the dots is used as the ground truth label. For the ABCD dataset, the gender of each subject was used as label to be predicted. In addition, for simulated ‘double moon’ and ABCD cortical thickness features, the support vector machine (SVM) with Gaussian kernel was used for cross-domain classification after harmonizing. For ABCD T1-weighted MRI data, a 3D CNN network was utilized for gender classification after harmonizing. As shown in Table 3 and Fig.6, the proposed MCD-GAN could guarantee the improvement of task performance after harmonization.

**Table 3.**
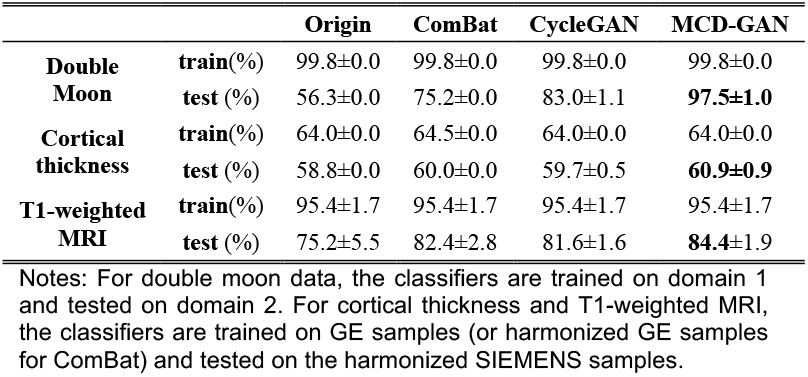
Comparison of classification performance of harmonization methods.

**Fig. 6.**
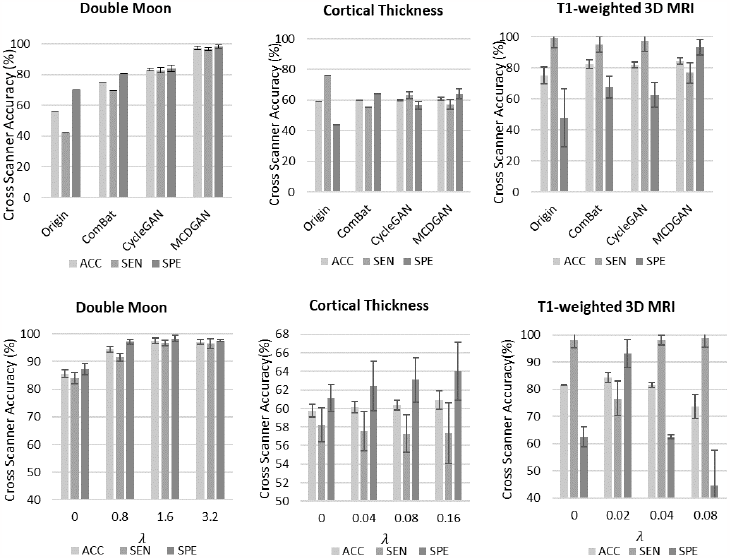
The comparison of downstream classification performances of different harmony approaches. (a) The comparison of the accuracy of cross-scanner classification; (b) The effects of hyper-parameter λ which controls the discrepancy ratio. For double moon data, the optimized λ is 1.6, and for cortical thickness feature, the optimized λ is 0.1. For T1-weighted 3D MRI, the optimized λ is around 0.02. Notes: ACC: accuracy; SEN: sensitivity; SPE: specificity.

For MCD-GAN, the hyperparameter *λ* influenced the data harmony effects by controlling the weight of classifier discrepancies. Here, we compared the downstream classification performance under different *λ*. when the *λ* was set to 0, the MCD-GAN degenerated to CycleGAN. The results are shown in Fig 6b. and Fig 7a. As *λ* increased exponentially, the classification performance increased, however, when *λ* was above a threshold, the classification performance started to decrease. For double moon data, the optimized *λ* was 1.6, for cortical thickness feature, the optimized *λ* was 0.1. For T1-weighted 3D MRI, the optimized *λ* was 0.02.

**Fig. 7.**
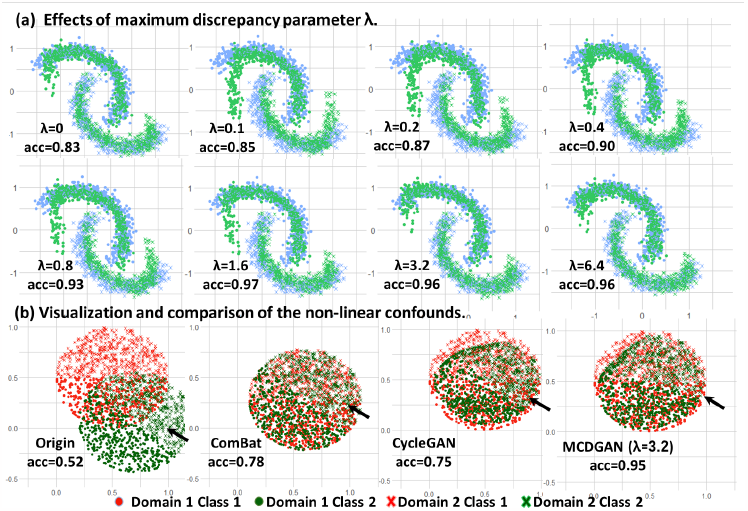
(a) Visualization of the effects of maximum discrepancy control hyperparameter *λ. λ* ∈ {0, 0.1,0.2,0.4,0.8,1.6,3.2,6.4}, ‘acc’ represents the cross-domain classification accuracy; (b) Visualization and comparison of harmonization methods for non-linear confounds. The simulated “double pancake” data has two domains: domain 1 contains 2D points that are randomly scattered in a circle. The sample size is 1000. The samples whose Y value over 0.5 is Category 1, and the samples whose Y value is below 0.5 is Category 2. Domain 2 points are generated by translation and rotating domain 1 by 30 degrees counterclockwise. MCD-GAN can thus correctly harmonize the non-linear confounds which are ignored by ComBat and CycleGAN (as arrow points). ‘acc’ represents the cross-domain classification accuracy.

## V. Discussion

Harmonizing scanner-related confounds is essential for improving cross-scanner reproducibility in multi-site neuroimaging studies. Conventional data harmony methods including residual-based and GAN-based are not designed to theoretically guarantee improved performance on specific downstream tasks. By leveraging the advantages of both data harmony and domain adaptation, our proposed MCD-GAN has three main advantages: ***1)*** MCD-GAN can theoretically guarantee improved performances on specific downstream tasks after harmonization; ***2)*** MCD-GAN reduces inter-scanner heterogeneity while preserving the anatomical layout of images; ***3)*** Different from ComBat which requires the change of all samples, the MCD-GAN harmonizes samples by mapping the target domain samples to the source domain, making it more flexible and suitable for continuous harmonizing.

Data harmony and domain adaptation are two categories of approaches for improving the consistency among multiple datasets and modalities. Data harmony approaches focus on estimating the distribution of the confounds and then utilizing specific algorithms to remove the confounds from original features, without breaking the anatomical information of original samples. Domain adaptation approaches focus on searching the domain invariant subspace. Therefore, if the distributions of confounds can be accurately estimated and are not relevant to the downstream tasks, data harmony should be preferred because the harmonized samples are general to various downstream tasks. However, when the confounds are usually non-linear and impractical to be accurately estimated, domain adaptation methods are preferred because the downstream tasks are considered. Our proposed MCD-GAN is a domain adaptation algorithm that preserves the anatomical information of original images, making it viable for neuroimaging studies.

The non-linear property of deep neural networks makes them unintuitive for interpretation, especially when input features are high-dimensional. To visualize how different harmony methods manipulate non-linear confounds, a simulated ‘double pancake’ was utilized. As shown in Fig. 7b, samples in domain 2 (green) were generated using domain 1 (red) samples by anti-clockwise 30° and then displacement. The ComBat only linearly transformed the domain 1 and domain 2 samples, ignoring the nonlinear confound, causing the failure of downstream classification after harmony. CycleGAN also failed to modify the distribution of generated samples according to the downstream classification task. Differently, our proposed MCD-GAN could improve the downstream classification performance by considering categorical information. Therefore, the classifiers trained on domain 1 can achieve satisfying performance when directly migrated to the harmonized data for conducting specific classification tasks.

Some open issues exist and should be addressed in future work. First, the proposed MCD-GAN utilizes CycleGAN for preserving the anatomical layout of the images. This means harmonizing datasets which contain *K* scanners requires training *K-1* models to map the samples from *K-1* domains into the selected domain one by one. As the *K* increases, the computational complexity would increase linearly. Thus, in the future, a balance between the computational complexity and the performance in removing confounds is to be studied. Second, in this work, we mainly focused on scanner effects, however, other confounds (e.g., age, race, educational years) may also hamper the reproducibility of neuroimaging studies. These need to be further studied in future work. Third, the GAN-based approach usually requires more samples for training in comparison to ComBat, it is worthy studying how to reduce training samples while maintaining the harmonization performance. Fourth, the proposed MCD-GAN model is a general framework that can also be extended by designing the classifiers according to specific downstream tasks, such as mental disorder classification, segmentation, and regression.

## VI. Conclusion

We propose the MCD-GAN, which takes advantage of both adversarial generative network and maximum discrepancy classifier approaches, for harmonizing the scanner effects, preserving the anatomical information, and improving the downstream cross-scanner classification performances. The advantage of the MCD-GAN was validated on simulated data and the ABCD MRI dataset. The mechanisms of MCD-GAN and the hyper-parameters were theoretically proved, visualized, and tested. In summary, MCD-GAN, as a general framework, is promising to facilitate cross-site reproducibility effectively in broad neuroimaging studies.

## References

[1] G. Varoquaux, “Cross-validation failure: Small sample sizes lead to large error bars,” NeuroImage, vol. 180, pp. 68-77, 2018/10/15/ 2018, doi: 10.1016/j.neuroimage.2017.06.061.

[2] N. K. Dinsdale, M. Jenkinson, and A. I. L. Namburete, “Deep learning-based unlearning of dataset bias for MRI harmonisation and confound removal,” Neuroimage, vol. 228, p. 117689, Mar 2021, doi: 10.1016/j.neuroimage.2020.117689.

[3] F. Hu et al., “Image harmonization: A review of statistical and deep learning methods for removing batch effects and evaluation metrics for effective harmonization,” NeuroImage, vol. 274, p. 120125, 2023/07/01/ 2023, doi: 10.1016/j.neuroimage.2023.120125.

[4] R. Garcia-Dias et al., “Neuroharmony: A new tool for harmonizing volumetric MRI data from unseen scanners,” NeuroImage, vol. 220, p. 117127, 2020/10/15/ 2020, doi: 10.1016/j.neuroimage.2020.117127.

[5] J.-P. Fortin et al., “Harmonization of cortical thickness measurements across scanners and sites,” NeuroImage, vol. 167, pp. 104-120, 2018/02/15/ 2018, doi: 10.1016/j.neuroimage.2017.11.024.

[6] V. M. Bashyam et al., “Medical Image Harmonization Using Deep Learning Based Canonical Mapping: Toward Robust and Generalizable Learning in Imaging,” arXiv preprint, vol. 1409.0473, 2020.

[7] J.-P. Fortin et al., “Harmonization of multi-site diffusion tensor imaging data,” Neuroimage, vol. 161, pp. 149–170, 2017.

[8] W. E. Johnson, C. Li, and A. Rabinovic, “Adjusting batch effects in microarray expression data using empirical Bayes methods,” Biostatistics, vol. 8, no. 1, pp. 118–127, 2007.

[9] G. Modanwal, A. Vellal, M. Buda, and M. A. Mazurowski, “MRI image harmonization using cycle-consistent generative adversarial network,” in Medical Imaging 2020: Computer-Aided Diagnosis, 2020, vol. 11314: International Society for Optics and Photonics, p. 1131413.

[10] V. M. Bashyam et al., “Deep Generative Medical Image Harmonization for Improving Cross-Site Generalization in Deep Learning Predictors,” Journal of Magnetic Resonance Imaging, 2021, doi: 10.1002/jmri.27908.

[11] H. Guan, Y. Liu, E. Yang, P.-T. Yap, D. Shen, and M. Liu, “Multi-site MRI harmonization via attention-guided deep domain adaptation for brain disorder identification,” Medical Image Analysis, vol. 71, p. 102076, 2021/07/01/ 2021, doi: 10.1016/j.media.2021.102076.

[12] J.-Y. Zhu, T. Park, P. Isola, and A. A. Efros, “Unpaired image-to-image translation using cycle-consistent adversarial networks,” in Proceedings of the IEEE international conference on computer vision, 2017, pp. 2223–2232.

[13] R. Wang, P. Chaudhari, and C. Davatzikos, “Embracing the disharmony in medical imaging: A Simple and effective framework for domain adaptation,” Med Image Anal, vol. 76, p. 102309, Nov 26 2021, doi: 10.1016/j.media.2021.102309.

[14] K. Saito, K. Watanabe, Y. Ushiku, and T. Harada, “Maximum classifier discrepancy for unsupervised domain adaptation,” in Proceedings of the IEEE conference on computer vision and pattern recognition, 2018, pp. 3723–3732.

[15] P. Julián Alberto, S. Diego Fernandez, and F. Enzo, “Unsupervised domain adaptation via CycleGAN for white matter hyperintensity segmentation in multicenter MR images,” in Proc.SPIE, 2020, vol. 11583, doi: 10.1117/12.2579548. [Online]. Available: 10.1117/12.2579548

[16] M. Wang and W. Deng, “Deep visual domain adaptation: A survey,” Neurocomputing, vol. 312, pp. 135–153, 2018, doi: 10.1016/j.neucom.2018.05.083.

[17] S. Ben-David, J. Blitzer, K. Crammer, A. Kulesza, F. Pereira, and J. W. Vaughan, “A theory of learning from different domains,” Machine Learning, vol. 79, no. 1, pp. 151-175, 2010/05/01 2010, doi: 10.1007/s10994-009-5152-4.

[18] S. Ben-David, J. Blitzer, K. Crammer, and F. J. A. i. n. i. p. s. Pereira, “Analysis of representations for domain adaptation,” vol. 19, p. 137, 2007.

[19] B. J. Casey et al., “The Adolescent Brain Cognitive Development (ABCD) study: Imaging acquisition across 21 sites,” Dev Cogn Neurosci, vol. 32, pp. 43–54, Aug 2018, doi: 10.1016/j.dcn.2018.03.001.

[20] D. J. Hagler, Jr. et al., “Image processing and analysis methods for the Adolescent Brain Cognitive Development Study,” Neuroimage, vol. 202, p. 116091, Nov 15 2019, doi: 10.1016/j.neuroimage.2019.116091.

